# Fast fronto-parietal cortical dynamics of conflict detection and context updating in a flanker task

**DOI:** 10.1101/336958

**Authors:** Christopher R. Brydges, Francisco Barceló, An T. Nguyen, Allison M. Fox

**Author notes:** Corresponding author: Christopher R. Brydges, School of Psychological Science (M304), University of Western Australia, 35 Stirling Highway, Perth, WA 6009.

## Abstract

The traditional N2 and P3 components of the human event-related potential (ERP) have been associated with cognitive control. However, recent research has found that the traditional target P3 consists of a mixture of stimulus-locked, response-locked, and latency variable P3-like positivities that can be functionally and topographically dissociated from one another. The current study aimed to extend previous results by examining target N2 and P3-like subcomponents indexing conflict detection and context updating at both low- and high-order levels in the neural hierarchy during control of target detection in a flanker task. Electroencephalographic (EEG) signals were recorded from 45 young adults while they completed a hybrid go/nogo flanker task, and Residue Iteration Decomposition (RIDE) was applied to functionally dissociate these peaks and establish their roles in conflict detection and context updating. Frequentist and Bayesian analyses showed a stimulus-locked frontal N2 revealing early conflict detection and fast categorization of nogo, congruous and incongruous trials, resulting in subsequent frontal P3-like activity (high-order context updating) elicited by infrequent nogo trials in the latency-variable RIDE cluster, and by the perceptually difficult incongruous trials in the response-locked cluster. In turn, the perceptually easier and more frequent congruous trials did not elicit frontal P3-like activity, although all trial types did elicit parietally distributed P3-like activity (low-order context updating), mostly within the latency-variable cluster. These novel findings support the presence of up to three distinct high-order task-set units regulating performance in our flanker task, as indexed by split-second, dynamic information transmission across frontal and parietal scalp regions.

## 1. Introduction

Cognitive control refers to a group of processes associated with performance of specific tasks through appropriate adjustments in executive attention and response selection, whilst minimizing interference from conflicting information (Botvinick, Braver, Barch, Carter, & Cohen, 2001; Botvinick, Cohen, & Carter, 2004), and is associated with neural activation across a widely distributed fronto-parietal cortical network for cognitive control (Niendam et al., 2012). Event-related potential (ERP) studies have consistently reported a series of peaks putatively contributed by this fronto-parietal network and associated with two temporarily contiguous higher-order cognitive processes: conflict detection and context updating. These two cognitive operations have been most notably associated with the frontal N2 (circa 200-350 ms post-stimulus onset) and the fronto-parietal P3 (circa 300-600 ms poststimulus onset) ERP components (Gratton, Cooper, Fabiani, Carter, & Karayanidis, 2018; Nguyen, Moyle, & Fox, 2016; van Veen & Carter, 2002). Recent ERP research in task switching, however, has suggested that the traditional division of the P3 complex into a frontal ‘novelty’ P3a (Friedman, Cycowicz, & Gaeta, 2001) and a centro-parietal ‘context updating’ P3b component (Donchin & Coles, 1988; Polich, 2007) may be overly simplistic. In turn, new evidence suggests that there may be multiple overlapping P3-like positivities putatively rising from activation across the fronto-parietal network, each with subtly distinct scalp topographies that are involved in handling rapidly changing cognitive demands (Barceló & Cooper, 2018; Brydges & Barceló, 2018; Enriquez-Geppert & Barceló, 2018). The aim of the current study was to functionally disentangle these fronto-parietal P3-like positivities, together with the temporally earlier frontal N2, and establish their roles in context updating and conflict detection, respectively.

### 1.1. N2 ERP Component

The frontal N2 has been described as a fronto-central negativity that is commonly examined in conflict detection and response inhibition tasks such as the go/nogo and the Eriksen flanker tasks (Nieuwenhuis, Yeung, & Cohen, 2004; Yeung & Nieuwenhuis, 2009). Its amplitude is typically enhanced on infrequently presented nogo and incongruent flanker trials compared to standard go and congruent flanker trials, and it is thought to reflect detection of conflict prompting for suppression of pre-potent responses and competing but inappropriate responses (Folstein & Van Petten, 2008). Although few studies have directly examined conflict detection during nogo and competing incongruous responses in flanker tasks, recent comparisons using a hybrid nogo/flanker task reported significantly delayed and more posterior N2 effects on incongruent flanker trials compared to nogo trials, suggesting differential engagement of conflict detection during these two cognitive control sub-processes (Brydges et al., 2012; Brydges, Anderson, Reid, & Fox, 2013). Additionally, previous fMRI and ERP source localization studies have implicated the anterior cingulate cortex in conflict detection and elicitation of the frontal N2 peak (e.g., Bekker, Kenemans, & Verbaten, 2005; Botvinick et al., 2004; Carter & van Veen, 2007; van Veen & Carter, 2002).

### 1.2. P3 ERP Component

Traditionally, the P3 component has been viewed as a dichotomous component, split into a frontal P3a that is associated with processing of surprising stimuli (Friedman et al., 2001), and a parietal P3b component that is thought to index ‘context updating’ operations (Donchin & Coles, 1988; Polich, 2007). That is, when the subject updates their model of (or belief about) the environment following a motivationally significant event, then the amplitude of the P3b peak is enhanced (Donchin & Coles, 1988; cf. Friston, FitzGerald, Rigoli, Schwartenbeck, & Pezzulo, 2017). Alternatively, Verleger (1997) reviewed the use of the P3 component in mental chronometry and found that its latency is partly, but not completely, associated with both stimulus encoding and behavioral response time, suggesting that this component is strictly locked neither to stimulus nor response onset. Additionally, Verleger (1997) found that response selection in complex choice tasks can result in a second, later P3-like waveform (see also Falkenstein, Hohnsbein, & Hoormann, 1994), providing evidence of a multiplicity of functionally distinct fronto-parietal P3-like positivities associated with cognitive control.

Recent task switching ERP research by Barceló and colleagues has also suggested the presence of multiple P3-like positivities overlying fronto-parietal scalp regions, further indicating that the traditional P3a/P3b dichotomy may be overly simplistic (Barceló & Cooper, 2018; Brydges & Barceló, 2018). Barceló and Cooper (2018) tested 31 young adults on a two-choice cued task-switching paradigm, a go/nogo task, and an oddball task, each using identical stimuli but requiring different cognitive and motor demands, and found that a target-locked P3 peak was evoked in all tasks and conditions, providing evidence for a domain general P3 component. Additionally, a large late positive complex (LPC) was also observed when a visual target (requiring a response) immediately followed a transition cue. Importantly, this LPC component was modulated by cognitive demands formalized as the amount of contextual information conveyed by the stimuli for subsequent response selection. That is, although the LPC was largest on target trials immediately following a switch cue, it was also elicited, albeit at a reduced amplitude, by target trials immediately following a repeat cue, and even after nogo trials in the go/nogo task. In turn, this LPC was absent several trials after the onset of switch/repeat cues, nogo events, and throughout the oddball task. These graded modulations in target P3-like activity tracked dynamic trial-by-trial changes in contextual information (see Fig. 1). Barceló and Cooper (2018) analyzed the current source densities of target P3 (300-350 ms) and target LPC (400-1100 ms) at electrode sites overlying fronto-parietal regions, and reported fast changing and subtle but significant differences in scalp topography, implying different configurations of fronto-parietal generators for different P3-like sub-components in first target trials. The fronto-parietal distribution of the P3/LPC complex has been linked to widespread activation across a fronto-parietal network (Bledowski et al., 2004), including the lateral prefrontal cortex, temporo-parietal junction, and pre-supplementary motor area (Niendam et al., 2012), consistent with a multiple demand system for cognitive control (Duncan, 2010).

**Figure 1.**
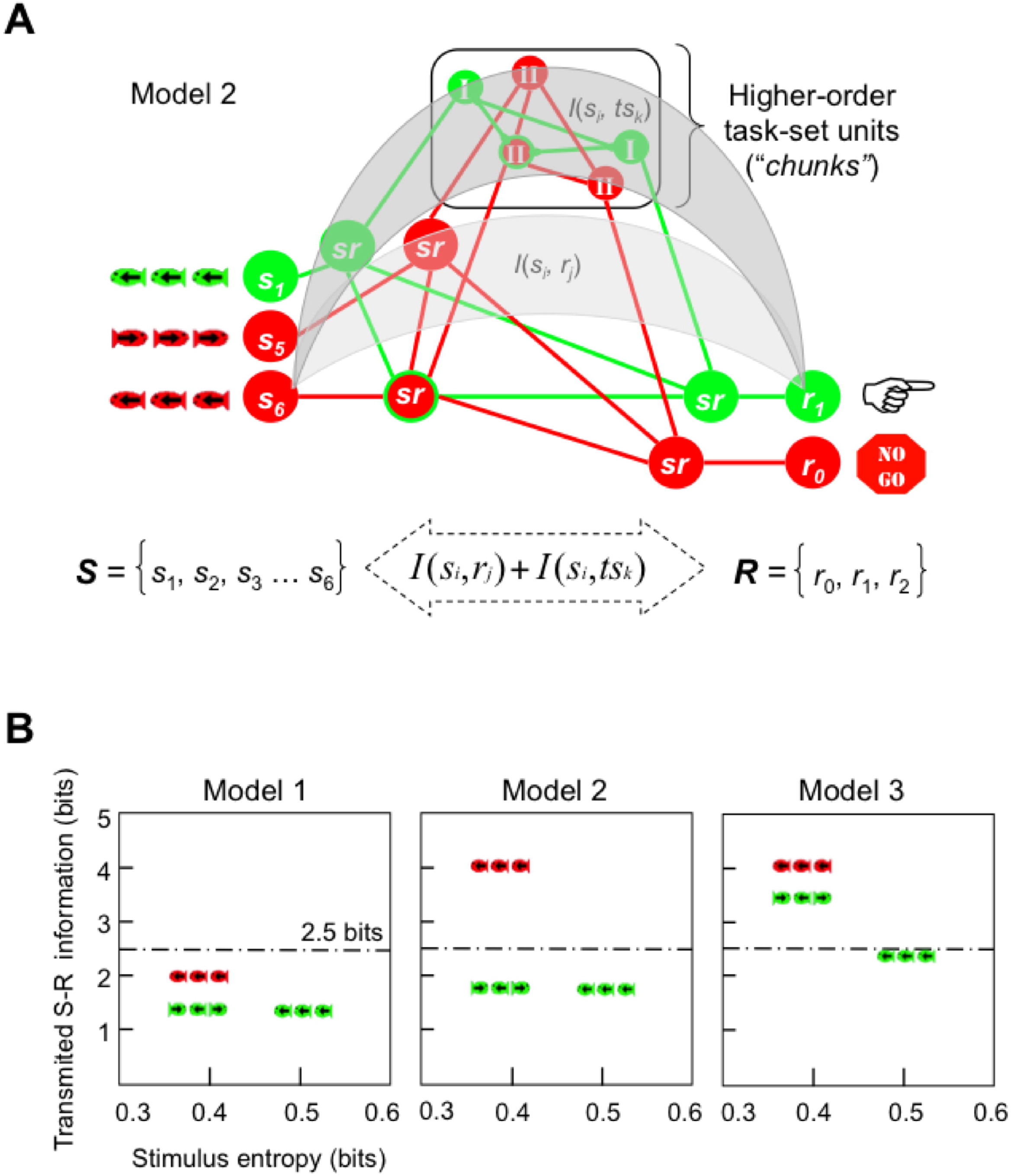
Formal modelling of cognitive demands in the flanker task. (A) An integrative model of prefrontal executive control (adapted from Miller & Cohen, 2001) was adopted to formalize contextual information in the flanker task in terms of all sensory, motor and intermediate low- and high-order sensorimotor representations (or task-set units), as well as their probabilistic interdependencies, inasmuch as an active working memory representation of all those elements was necessary for efficient flanker task performance. For simplicity, only three trial types are illustrated here, namely, a congruous row of green left-facing fish is mapped onto a left-hand response, whereas two rows of red fish are mapped onto a nogo response. Actual stimulus displays consisted of five green or red fish in a row, and the full stimulus set consisted on six stimulus exemplars that could be mapped either to left-hand, right-hand, or nogo responses. Cognitive demands were estimated in terms of sensorimotor information transmission across both low- and high-order levels within the putative hierarchy of cognitive control. (B) A priori estimations of transmitted information, *I*(*s_i_*, *r_j_*)+ *I*(*s_i_*, *tS_k_*), between stimuli and responses as a function of stimulus entropy, *H*(*s_i_*)= −*p*(*s_i_*)· log_2_ *p*(*s_i_*), of congruous, incongruous and nogo stimuli. The information transmitted from stimuli to responses is derived from the notion of mutual information, *I*(*S*; *R*), between the set of all stimuli, *S* {*s*_1_, *s*_2_, *s*_3_, *s*_4_, *s*_5_, *s*_6_}, and their associated responses, *R* {*r*_0_ *r*_1_, *r*_2_} (cf. Attneave, 1959; Koechlin & Summerfield, 2007; see details in Supplementary material). The dotted line marks the theoretical human capacity for holding information in working memory: 2.5 bits (cf. Miller, 1956). Accordingly, the same trial type could convey different amounts of transmitted information depending on the hypothetical number of hidden high-order task set units (or memory “chunks”) assumed to regulate performance in the flanker task. Model 1: 1 chunk, model 2: 2 chunks; model 3: 3 chunks (see the main text for a full explanation).

One limitation of traditional ERP analyses to examine neurophysiological processes is that they are typically locked to a particular task event – generally either a sensory stimulus or a motor response (Luck, 2014) – when both stimulus-locked and response-locked activity are elicited within the same time epoch, and hence, they potentially temporally overlap each other. As a consequence, an ERP component that is locked to trial-variable reaction times (RTs) will not show a clear peak and will be smeared within the stimulus-locked ERP waveform, and vice versa. In an attempt to counter this, Ouyang and colleagues (Ouyang, Herzmann, Zhou, & Sommer, 2011; Ouyang, Hildebrandt, Sommer, & Zhou, 2017; Ouyang, Sommer, & Zhou, 2015, 2016) developed a novel method for separating ERP components based on the variability of single trial latencies, namely residue iteration decomposition (RIDE). This technique defines clusters of components that are either stimulus-locked (S cluster), response-locked (R cluster), or neither (i.e., affected partially, but not completely, by RT – referred to henceforth as the central C cluster). The RIDE technique is implemented by iteratively calculating correlations between estimates of stimulus-locked and response-locked components and EEG waveforms across single trials (Ouyang et al., 2011).

Brydges and Barceló (2018) applied RIDE analyses to EEG data on the same two-choice cued switching task, and showed a variety of functionally distinct albeit topographically similar P3-like positivities across fronto-parietal regions for each of the stimulus-locked, response-locked, and latency-variable (central) clusters, each with subtly distinct amplitude and topographical modulations occurring as a function of trial-by-trial demands. Specifically, regular P3-like (300-350 ms) peaks were observed in all three clusters and were modulated by task-specific contextual information, implying that the context updating hypothesis of P300 elicitation (Donchin & Coles, 1988) may be reconceptualized in terms of trial-by-trial updating of the mental model of the task (aka the “task-set”) putatively being held at fronto-parietal networks (Friston et al., 2017). A corollary of this new proposal is that context updating can be triggered by adjustments at various levels in the putative hierarchy of cognitive control (Brydges & Barceló, 2018; Miller & Cohen, 2001), either due to changes in sensory input, changes in motor responses, and/or changes in low- and high-order intermediate sensorimotor processes (Fig. 1A). Additionally, some of these frontoparietal P3-like positivities showed a more frontal scalp distribution under some specific trial conditions, thus reflecting rapid increases in frontal resources dynamically engaged to meet trial-by-trial changes in cognitive demands (Koechlin & Summerfield, 2007). In sum, these results suggest that the target P3 complex consists of several functionally and topographically distinct stimulus-locked, intermediate latency-variable, and response-locked P3-like positivities contributed by a widely distributed fronto-parietal network (Niendam et al., 2012). This proposed P3 taxonomy implies multiple functionally and temporally distinct, though partly overlapping context updating operations that can be extracted from the traditional ERP waveform.

### 1.3. Miller and Cohen’s (2001) Theory of Prefrontal Function

Miller and Cohen (2001) posited that “cognitive control stems from the active maintenance of patterns of activity in the prefrontal cortex that represent goals and the means to achieve them” (p. 167). Accordingly, one function of the prefrontal cortex is to supervise and coordinate signals to and between other neural regions that are responsible for encoding incoming information (e.g., a stimulus), applying various rules to the information (e.g., response conflict resolution), and the resulting output (e.g., selecting an appropriate response). This coordination aspect is especially important when presented stimuli are ambiguous (i.e., activate more than one input representation) or when there is response conflict. The schematic representation shown in Figure 1A illustrates these postulated operations, highlighting the sensory processing units indicative of neural representations of the sensory event together with response processing units indicative of the selection of the appropriate response options. The role of the prefrontal cortex is to activate and coordinate neural pathways to ensure the selection of the appropriate response. The stimuli from the hybrid flanker go/nogo task used in the current study on the left, the response options are on the right, and in the middle are hidden task set units (though in order to keep the model relatively simple, only three of six possible stimulus types are shown, including no incongruous stimuli, and only two out of three possible response options are shown). In the case of Figure 1A, the role of the prefrontal cortex is to activate and coordinate neural pathways so that presentation of a red nogo stimulus results in no response being made. From this view, the prefrontal cortex guides the flow of neural activity to update the stimulus-response mapping rule for the current nogo trial (i.e., no response is to be made), then to the neural region associated with inhibiting a motor response. From there, these pathways can be strengthened through repeated practice and successful learning. The multi-level structure of this model of prefrontal function is compatible with the multiple and functionally distinct framework of updating processes posited by the updated P3 hypothesis.

### 1.4. The Current Study

The current study aimed to extend previous results by examining target N2 and P3-like subcomponents indexing conflict detection and context-updating processes at both low- and high-order levels in the neural hierarchy during cognitive control in a flanker task. In doing so, we adopted a widely cited model of prefrontal executive control (Miller & Cohen, 2001; Fig. 1A), together with formal information theory estimates of cognitive demands associated with the processing of nogo, congruent, and incongruent flanker trials (Cooper, Darriba, Karayanidis, & Barceló, 2016; Koechlin & Summerfield, 2007; Miller, 1956; Fig. 1B). This formal framework helped us characterize the relative contribution from between-condition differences in sensory, motor and intermediate sensorimotor task-set units towards the updating of the ongoing task context, as the participants’ uncertainty about any of the changes was assumed to elicit conflict detection (frontal N2) and context updating (frontoparietal P3) operations (Barceló & Cooper, 2018; Donchin & Coles, 1988; Friston et al., 2017). The task context was defined as any sensory, motor, or intermediate sensorimotor neural representations (also high-order task-set units, or memory “chunks”; Fig. 1A), as well as their probabilistic interdependencies (cf. Friston et al., 2017), inasmuch as an active working memory representation of all those elements is necessary for efficient task performance. The task consisted of a hybrid go/nogo flanker task (Brydges et al., 2012, 2013) where a congruous row of green fish was to be categorized using left or right button presses according to the direction of the central fish. In a small proportion of trials, the direction of the central fish was incongruent to that of surrounding flanker fish. Further, a small proportion of ‘nogo’ trials consisted of congruous fish presented in a different color (red). Through explicit task instructions and practice, participants could quickly acquire the correct stimulus-response (S-R) mappings to be implemented through low-order sensorimotor S-R links (Fig. 1A). Importantly, the dynamic engagement of prefrontal executive resources on a trial-by-trial basis was hypothesized to depend on the number of ‘hidden’ high-order task-set units necessary for regulating the dynamic updating of the same constant number of low-order sensorimotor units at posterior association cortices, with a larger number of high-order task-set units requiring larger prefrontal resources. The dynamic trial-by-trial changes in cognitive demands associated with the processing of different flanker stimuli could thus be modelled as information transmission through hypothetical (hidden) high-order task-set units (Figs. 1A,B; cf. Friston et al., 2017), with one high-order task-set unit implying no transmission of information through prefrontal cortices (model 1; Fig. 1B), and progressively larger numbers of high-order task-set units involving gradually larger amounts of information transmission through prefrontal regions (Fig. 1B; Table 1; see Supplementary materials). The feasibility of each of these models was then assessed using Bayesian model comparison on the electrophysiological evidence provided by RIDE decomposed target N2 and P3-like activity from the C, S and R clusters, and specifically the activity recorded over frontal scalp regions.

**Table 1.** Modeling of transmitted S-R information (in bits) for the three task conditions in Fig. 1, as a function of the number of hypothetical high-order task-set units (or memory “chunks “)

| Task Conditions | Model 1<br>(1 "chunk") | Model 2<br>(2 "chunks") | Model 3<br>(3 "chunks") |
| --- | --- | --- | --- |
| *Congruous* | 1.42 | 1.84 | 2.42 |
| *Incongruous* | 1.42 | 1.84 | 3.42 |
| *NoGo* | 2.00 | 4.00 | 4.00 |
| Low-order $\Sigma I(s_i, r_j)$ | 4.84 | 4.84 | 4.84 |
| High-order $\Sigma I(s_i, ts_k)$ | 0.00 | 2.84 | 5.00 |

Note. Models considering different high-order task-set units for left and right stimuli and responses were dismissed as implausible, given evidence of similar target N2 and P3 responses irrespective of the left versus right direction of flanker stimuli. For technical details, see Supplementary materials.

Under the assumption that the frontal N2 component is elicited as a result of conflict detection (Folstein & van Petten, 2008; Nguyen et al., 2016; van Veen & Carter, 2002), whereas the target P3 is elicited as a result of context updating operations (Barceló & Cooper, 2018; Donchin & Coles, 1988; Friston et al., 2017), several hypotheses can be derived from the theoretical rationale described above, depending on the number of high-order task-set units involved in regulating efficient flanker task behavior. For instance, if only one high-order task-set unit was needed (model 1), then electrophysiological differences between task conditions should be apparent over parietal scalp sites only. Alternatively, our model’s predictions differ depending on whether two high-order task-set units (i.e., one to categorize the frequent green targets, and one to categorize the infrequent red nogo trials; model 2), or three units were involved (i.e., one for frequent congruous targets, one for infrequent incongruous targets, and one for infrequent nogo trials; model 3, Fig. 1B), with N2-P3 effects at frontal regions predicted only for those task conditions that overshoot working memory capacity (reflected by the 2.5 bit limit in each model in Fig. 1B; cf. Miller, 1956). These models assume the same amount of transmitted information through low-level sensorimotor units for the six S-R links required to perform the task, and hence, the main differences among models depend on the number of hidden high-order task-set units in the hierarchy of control (Fig. 1A; Table 1). As such, and based on the extant literature, it was expected that conflict detection (reflected in the frontal N2 peak) would be a prerequisite for context updating (target P3 and LPC). Additionally, based on Brydges and Barceló’s (2018) revised version of the context updating theory, it was hypothesized that the target P3 complex would be elicited by multiple context updating operations, namely, by the updating of sensory units (as evidenced by observing between-conditions differences in P3-like positivities in the S cluster), the updating of motor units (R cluster), or the updating of intermediate sensorimotor units (C cluster). Further, the scalp topographies of these positivities were expected to differ subtly between task conditions, with more frontally distributed scalp topographies for the most informative flanker stimuli (Fig. 1B, Table 1). Based on the earlier latency of the frontal target N2 component, and its role in conflict detection, task effects were mostly predicted for the stimulus-locked cluster, as a prerequisite stage for subsequent context updating indexed by target P3-like activity.

## 2. Method

### 2.1. Participants

The current study combined the adult datasets of two previous studies (*n* = 12 from Brydges et al., 2012; *n* = 13 from Brydges et al., 2013), plus 20 previously unpublished datasets. In total, 45 undergraduate psychology students (*M*_age_ = 21.93 years, *SD*_age_ = 6.30 years, range = 18-49 years); 26 females and 19 males; 37 right-handed and 8 left-handed) were recruited for the study. All participants self-reported normal or corrected-to-normal vision, no history of colour blindness or neurological disorders, and each participant performed above chance on all three task conditions. The samples were combined in order to increase statistical power given that neuroscientific studies are commonly underpowered (Button et al., 2013). The current hypotheses were only tested on the entire sample and not on the previously published samples.

### 2.2. Materials

The same hybrid go/nogo flanker task used by Brydges et al. (2012; 2013) was used in this study. Each stimulus consisted of five fish presented on a blue background. An arrow on the body of the fish specified direction and the target was the central fish. Participants were instructed to press a response button on a keyboard (red felt patches on the ‘Z’ and ‘/’ keys of a QWERTY keyboard) analogous to the direction of the central fish. The task had three conditions: in the congruent condition (.5 probability), the fish were green and all facing the same direction (.25 probability for left and right facing green fish, respectively). In the incongruent condition (.25 probability), the fish were also green, however the flankers faced the opposite direction to the central target (with .125 probability of left and right incongruent flankers, respectively). In the nogo condition (.25 probability), the fish were congruent and red (again with .125 probability of either left or right facing red fish), and participants were required to withhold their response. Each fish subtended .9° horizontally and .6° vertically, with .2° separating each fish (see Figure 2). A fixation cross was presented in the centre of the computer screen for 500 ms before the stimulus appeared immediately above it. Stimuli were presented in random order for 300 ms with a 2,000 ms inter-stimulus interval. The task was presented to the participants as a game in which they should feed the central fish. Speed and accuracy were equally emphasized. Eight practice trials were administered to ensure the participants understood the task requirements. A total of 176 trials were subsequently presented in one block.

**Figure 2.**
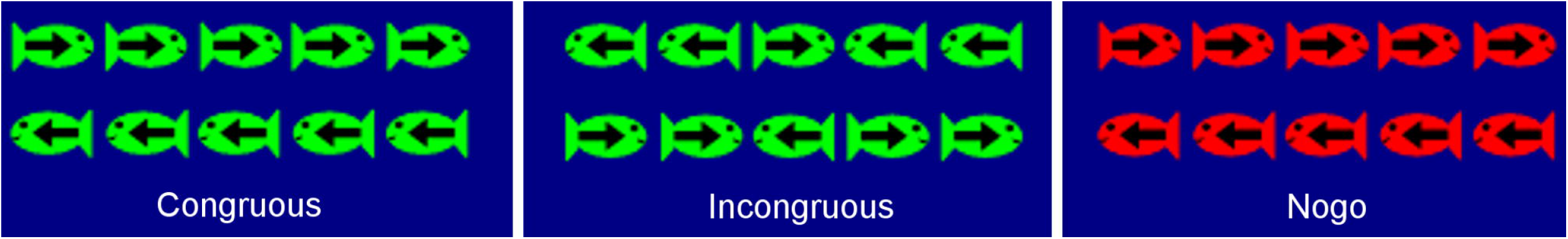
The six stimuli used in the current study.

### 2.3. Information Theoretical Estimations

We adopted an information theoretical model of cognitive control as a formal tool to help us operationalize the task set (or context) in terms of low- and high-order sensorimotor (S-R) information transmission within a putative hierarchy of fronto-parietal control processes (Fig. 1A; cf. Barceló & Knight, 2007; Barceló & Cooper, 2018). In doing so, we followed recommendations by Miller (1956) for estimating the amount of information transmitted between contextually related stimuli and responses (or *input-output correlations*). These estimates allowed us to define the informational structure of our flanker task in terms of, not only mean stimulus probabilities, but also joint probabilities among stimuli, their associated motor responses, and any relevant cognitive control operation putatively involved (e.g., updating of high-order task-set units; Fig. 1A). Thus, the task context was modelled at two hierarchically distinct levels: (1) low-order sensorimotor S-R links, and (2) hypothetical (hidden) high-order task-set units (Friston et al., 2017). Accordingly, the working memory representation of congruous green flanker targets and their associated responses (i.e., the low-order *s*_i_-*r*_j_ links in Fig. 1A) was to be twice as frequently activated compared to the working memory representation of incongruous green flanker targets, or that of red nogo distracters. Importantly, the dynamic engagement of prefrontal resources on a trial-by-trial basis was hypothesized to depend on the number of ‘hidden’ high-order task-set units necessary for regulating the dynamic updating of the six low-order sensorimotor units being held at posterior association cortices, with a larger number of high-order task-set units involving greater allocation of prefrontal resources (Fig. 1B; cf. Friston et al., 2017). Thus, whereas one high-order task-set unit implies no transmission of information through prefrontal cortices, three high-order task-set units involve an averaged transmission of 5.0 bits of information (Table 1). Models considering different high-order task-set units for left and right stimuli and responses were dismissed as implausible, given evidence of similar target N2 and P3 responses irrespective of the left versus right direction of flanker stimuli. Note that these information estimates can be seen as a more formal and accurate way to translate into bits the mean probabilities of task events, as is common practice in most experimental psychology studies. Yet they provide a common metric to compare different task conditions at both low- and high-order levels in the putative hierarchy of cognitive control. For instance, instead of saying that a left green congruous target occurs with an overall mean probability of *p* = .25 in our flanker task, we chose to quantify this in bits by saying that the sensory entropy of this trial type is: *H*(s_1_) = −0.25 · log_2_ 0.25 = 0.50 bits. A similar formalism was used to quantify in bits the relative probabilities of the six specific low-order *s*_i_-*r*_j_ links, and the hidden high-order *s_i_*-*ts_k_* links, using the concept of transmitted information: *I*(*s*_i_, *r*_j_)= log_2_ *p*(*s*_i_, *r*_j_) −log_2_ *p*(*s*_i_) −log2 *p*(*r*_j_) and *I*(*s*_i_, *ts*_k_)= log_2_ *p*(*s*_i_, *tS*_k_) −log_2_ *p*(*s*_i_) −log_2_ *p*(*tS_k_*) for low- and high-order levels in the hierarchy of control, respectively (Fig. 1A; cf. Miller, 1956). Table 1 offers a summary of these information-theoretic estimations; for a more detailed technical description, see the Supplementary material (cf. Barceló & Cooper, 2018).

### 2.4. Electrophysiological Acquisition

The EEG was continuously recorded using an Easy-Cap™ Ag/AgCl sintered ring electrodes were placed at 33 sites based on Easy-Cap montage 24. Eye movements were measured with bipolar leads placed above and below the left eye. The EEG was amplified with a NuAmps 40-channel amplifier, and digitized at a sampling rate of 250 Hz. Impedances were below 5 kΩ prior to recording. During recording, the ground lead was located at AFz and the right mastoid was set as reference.

EEG data were processed using MATLAB (Mathworks, Navick, MA) through a pipeline utilizing EEGLAB version 14.0.0 (Delorme & Makeig, 2004), ERPLAB version 6.1.3 (Lopez-Calderon & Luck, 2014) and ADJUST version 1.1.1 (Mognon, Jovicich, Bruzzone, & Buiatti, 2011). Preprocessing was performed in EEGLAB by re-referencing offline to a common average, and bandpass filtering the data (0.1 – 30 Hz). Epochs for each stimulus type were extracted from 100 ms prestimulus to 1,000 ms poststimulus onset.

Independent components analysis was conducted using the extended infomax algorithm (Bell & Sejnowski, 1995), and ADJUST was used to detect any artefactual components (including blinks, eye movements and muscle movement). These components were removed, and the remaining components were back-projected to the electrode space. The mean number of components removed per participant was 6.67 (*SD* = 3.36). Epochs containing EEG signals exceeding ±100 μV at any electrode site were excluded from analyses.

### 2.5. RIDE

RIDE analysis followed the methods described in Ouyang et al. (2011, 2015). The RIDE toolbox and manual can be found at http://cns.hkbu.edu.hk/RIDE.htm. RIDE decomposes ERPs into stimulus-locked, response-locked, and central clusters (terms S, R, and C, respectively, from here), though for the nogo condition, no R cluster was extracted as a correct nogo trial requires no response to be made (Ouyang, Schacht, Zhou, & Sommer, 2013). The latency estimates of S and R (L_S_ and L_R_) are the stimulus onset and response time, respectively. The latency estimate of C (L_C_) is derived from the data of each individual participant using the following iterative process.

#### 2.5.1. Initial estimation of L_C_

RIDE separates component clusters by estimating the latency of the S, C, and R clusters on single trials. It is assumed that the C cluster is not stimulus or response-locked, and that L_C_ is variable over single trials as a result of this. Based on L_S_, L_R_, and L_C_, the evoked potentials of S, R, and C are dissociated from one another in later steps. The decomposition module makes use of both external time markers (e.g., stimulus and response onset) and estimated component latencies. The latency-locked clusters (i.e., S and R clusters) are removed from the single-trial data before the latency of the non-locked latency-variable component cluster (the C cluster) is estimated, before the C and R clusters are removed to estimate the S cluster, then the S and C clusters are removed to estimate the R cluster. This process is then repeated in iterative manner, following the procedures described by Ouyang et al. (2011; 2015). An initial estimation of L_C_ was made with the Woody (1967) filter for each recording site between 250 and 550 ms. The mean latency across channels was taken as L_C_ across the scalp.

### 2.6. Statistical analyses

For both traditional ERP and RIDE analyses, 3 × 2 repeated measures ANOVAs were conducted on the data to identify mean differences in amplitude between conditions (congruous, incongruous, and nogo) and electrode site (Fz and Pz). These electrode sites were chosen based on Brydges and Barceló (2018). For the ERP analyses, mean amplitudes were analyzed between 220-270 ms (corresponding to the N2 component), 300-400 ms (P3), 450-550 ms (LPC_1_), and 600-700 ms (LPC_2_). These latency windows were chosen based on previous N2 and target P3 research (e.g., Barceló & Cooper, 2018; Falkenstein, Hohnsbein, & Hoormann, 1999) and visual inspection of the grand mean ERP waveforms. To ensure consistency, these same time windows were used in the RIDE analyses: the 220-270 ms and 300-400 ms were extracted from the S cluster, and the 300-400 ms, 450-550 ms, and 600-700 ms windows were extracted from the C cluster. Additionally, the R cluster was defined as the mean amplitude occurring 50 ms before a correct response was made, to 50 ms after the response. As mentioned previously, no R cluster was extracted for the nogo condition as a correct trial requires no response to be made, and hence, the statistical design was a 2 (site) × 2 (condition) repeated measures ANOVA. Additionally, a repeated measures ANOVA with condition as the only factor was conducted on the behavioral accuracy data, and a paired-samples *t*-test was conducted on the mean RTs for the congruous and incongruous conditions.

In addition to traditional null hypothesis significance testing, we conducted Bayesian model comparison (BMC; Wagenmakers, 2007; Wagenmakers et al., 2018). Bayesian statistics are advantageous over frequentist statistics for various reasons: first, BMC testing allows a hypothesis to be accepted or rejected by gathering evidence in favor of it (Dienes, 2011; Kruschke, 2013). That is, the alternative hypothesis can only be falsified by accepting the null hypothesis over it, which Bayesian statistics allow for. Second, Bayesian statistics allow the same data points to be repeatedly tested without researchers having to pre-commit to a specified sample size, whereas this cannot be done with frequentist statistics (Armitage, McPherson, & Rowe, 1969). Third, Bayesian statistics are produced in terms of the probability of hypotheses given data, as opposed to data given hypotheses (Cohen, 1994). Bayesian statistics are more interpretable than frequentist statistics (Dienes, 2011; Kruschke, 2013), and assess the credibility of one hypothesis compared to another. Hence, Bayesian methods are well-suited for testing hypotheses about which of the three models presented in Table 1 and Fig. 1B best explains ERP and RIDE data at frontal regions, particularly in order to counteract potential Type I errors associated with *p* values of conventional frequentist statistics (Luck & Gaspelin, 2017). A Bayes Factor (BF) was calculated from the Bayesian ANOVAs to test how much the data supported the alternative (H1) over the null (H0) hypothesis (that is, how strong the evidence was in favor of one model over another). Based on guidelines set by Jeffreys (1961), a BF_10_ > 3 was considered sufficient evidence in favor of the alternative hypothesis, and a BF_10_ > 10 was considered to be strong evidence in favor. Of note, BF_10_ refers to the BF value of H1 being supported over H0, whereas BF01 refers to the opposite. To calculate BF_01_, one simply inverts the BF_10_ value. Additionally, posterior probabilities (henceforth referred to as *p*(H|D)) evaluated the probability of a hypothesis being correct given the observed data, with values of .50-.75, .75-.95, .95-.99, and > .99 indicating weak, positive, strong, and very strong evidence in favor of the alternate hypothesis, respectively (Masson, 2011; Raferty, 1995). Alternative hypotheses would only be accepted if BF_10_ > 3. Both the frequentist and Bayesian analyses were conducted using JASP version 0.8.5.1 (JASP Team, 2018). To save space, simple effects of the frequentist ANOVA for Fz are only reported when they differ from the expected BMC results.

## 3. Results

### 3.1. Behavioral results

Descriptive statistics of the behavioral data are presented in Table 2. A repeated measures ANOVA found a main effect of condition on accuracy (*F*(2,88) = 72.98, *p* < .001, η_p_^2^ = .62), and subsequent post-hoc paired-samples *t*-tests found lower accuracy on the incongruous condition than the congruous and nogo conditions (congruous-incongruous *t*(44) = 9.83, *p* < .001, Cohen’s *d* = 1.99; congruous-nogo *t*(44) = 1.59, *p* = .12, Cohen’s *d* = 0.26; nogo-incongruous *t*(44) = 8.44, *p* < .001, Cohen’s *d* = 1.37). Also, mean RTs were longer in the incongruous condition compared to the congruous condition (*t*(44) = 15.80, *p* < .001, Cohen’s *d* = 1.19).

**Table 2.** Descriptive Statistics of Behavioral Results (means, with standard deviations in parentheses)

| Condition | Reaction Time (ms) | Accuracy (proportion correct) |
| --- | --- | --- |
| Congruous | 378 (43) | .98 (.03) |
| Incongruous | 447 (60) | .82 (.11) |
| Nogo | N/A | .96 (.06) |

### 3.2. Traditional ERP analyses

Figure 3 shows the grand average ERP waveforms for the three conditions at Fz and Pz sites, and mean amplitude scalp maps corresponding to five time windows in the recording epoch, corresponding to the N2, P3, LPC_1_, and LPC_2_ components. To aid comparison between traditional ERP components and RIDE clusters, Table 3 presents a summary of the main effects and interactions for the site and condition factors; however, for the sake of brevity, only those interactions, main effects, and post-hoc tests that are of theoretical importance are described in-text below.

**Figure 3.**
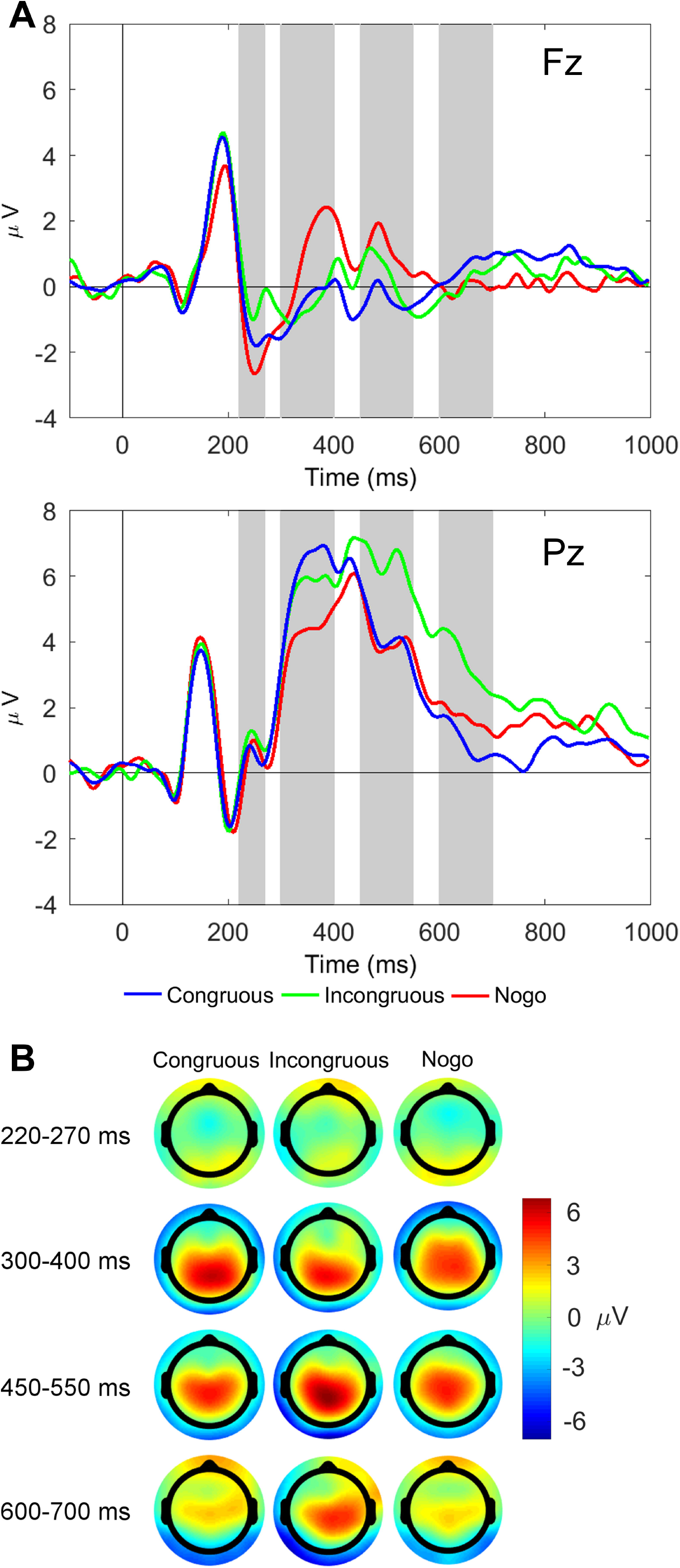
Stimulus-locked grand average ERP waveforms and scalp topography maps. (A) Waveforms depict mean voltages recorded from Fz (top) and Pz (bottom). Shaded areas indicate time windows used to measure mean ERP amplitudes tracking the temporal dynamics of the N2 and late P3-like complex: N2 (220-270 ms), P3 (300-400 ms), LPC_1_ (450-550 ms), and LPC_2_ (600-700). (B) Scalp topographies of the N2 and three late P3-like positivities depicted in (A) across task conditions (i.e., congruous, incongruous, and nogo).

**Table 3.** Summary of ANOVA results showing task effects for conventional ERP components and the RIDE decomposed C, S and R clusters.

|  | Site |  | Condition |  | Site x Condition |  |
| --- | --- | --- | --- | --- | --- | --- |
|  | *df* 1, 44 |  | *df* 1, 44 |  | *df* 2, 88 |  |
| ERPs | *F* | $\eta^2$ | *F* | $\eta^2$ | *F* | $\eta^2$ |
| N2 (220-270 ms) | 6.52 * | .13 | 8.75 *** | .17 | 2.30 *ns* | .05 |
| P3 (300-400 ms) | 55.35 *** | .56 | 0.72 *ns* | .02 | 21.35 *** | .33 |
| LPC<sub>1</sub> (450-550 ms) | 61.44 *** | .58 | 13.23 *** | .23 | 9.90 *** | .18 |
| LPC<sub>2</sub> (600-700 ms) | 7.72 ** | .15 | 4.23 * | .09 | 8.81 *** | .17 |
| RIDE S cluster | *F* | $\eta^2$ | *F* | $\eta^2$ | *F* | $\eta^2$ |
| sN2 | 0.99 *ns* | .02 | 10.81 *** | .20 | 5.76 ** | .12 |
| sP3 | 6.56 * | .13 | 3.98 * | .08 | 3.34 * | .07 |
| RIDE C cluster | *F* | $\eta^2$ | *F* | $\eta^2$ | *F* | $\eta^2$ |
| cP3 | 41.24 *** | .48 | 6.78 ** | .13 | 1.96 *ns* | .04 |
| cLPC<sub>1</sub> | 36.20 *** | .45 | 10.76 *** | .20 | 1.05 *ns* | .02 |
| cLPC<sub>2</sub> | 7.33 ** | .14 | 3.59 * | .08 | 2.67 *ns* | .06 |
| RIDE R cluster | *F* | $\eta^2$ | *F* | $\eta^2$ | *F* | $\eta^2$ |
| rP3 | 83.19 *** | .65 | 1.54 *ns* | .03 | 6.33 * | .13 |

Note. *ns*, non-significant, \* $p < .05$ , \*\* $p < .01$ , \*\*\* $p < .001$ . The *df* values for the R cluster interaction were 1, 44.

#### 3.2.1. N2 (220-270 ms)

The site × condition interaction was not significant, whereas the main effects of condition and site both reached significance (see Table 3). BMC testing at Fz found evidence in favor of a difference between conditions (B_F10_ = 93.50, *p*(H|D) = .99), with post-hoc tests finding evidence in favor of a smaller (less negative) incongruous N2 than the congruous (BF_10_ = 9.51) and nogo conditions (BF_10_ = 27.87), which did not differ between each other (BF_10_ = 0.88).

#### 3.2.2. P3 (300-400 ms)

The site × condition interaction and the main effect of site were both significant, whereas the main effect of condition was not (Table 3). Post-hoc t-tests found larger mean P3 amplitudes in the nogo condition than in the congruous and incongruous conditions at Fz (*t*(44) = 4.24, *p* < .001, and *t*(44) = 3.14, *p* = .003, respectively), with the reverse true at Pz (*t*(44) = −5.26, *p* < .001, and *t*(44) = −3.25, *p* = .002, respectively). BMC testing at Fz found evidence in favor of a difference between conditions (BF_10_ = 177.06, *p*(H|D) > .99), with the nogo P3 being larger than the congruous (BF_10_ = 211.36) and incongruous conditions (BF_10_ = 11.08), which did not differ between them (BF_10_ = 0.16).

#### 3.2.3. LPC1 (450-550 ms)

The site × condition interaction was significant, as were the main effects of site and condition (Table 3). Paired-samples t-tests found that the incongruous and nogo conditions elicited larger LPC_1_ amplitudes than the congruous condition at Fz (*t*(44) = 2.25, *p* = .029, and *t*(44) = 3.93, *p* < .001, respectively). At Pz, however, the incongruous condition evoked larger LPC_1_ amplitudes than the congruous and nogo conditions (*t*(44) = 4.81, *p* < .001, and *t*(44) = 4.30, *p* < .001, respectively). BMC testing at Fz found evidence in favor of a difference between conditions (BF_10_ = 20.33, *p*(H|D) = .95), with the nogo LPC_1_ being larger than the congruous (BF_10_ = 87.24). The evidence for/against differences between congruous and incongruous LPC_1_, and incongruous and nogo LPC_1_ was weak (BF_10_ = 1.58 and BF_10_ = 0.52, respectively).

#### 3.2.4. LPC2 (600-700 ms)

The site × condition interaction was significant, as were the main effects of site and condition (Table 3). Paired-samples t-tests found that the incongruous condition evoked larger mean LPC_2_ amplitudes than both the congruous and nogo conditions at Pz (*t*(44) = 6.05, *p* < .001, and *t*(44) = 2.45, *p* = .018, respectively). BMC testing at Fz found evidence against a difference in LPC2 between conditions (BF_10_ = 0.16, *p*(H|D) = .14). *Figure 3*. (1.5 columns) Stimulus-locked grand average ERP waveforms and scalp topography maps. (A) Waveforms depict mean voltages recorded from Fz (top) and Pz (bottom). Shaded areas indicate time windows used to measure mean ERP amplitudes tracking the temporal dynamics of the N2 and late P3-like complex: N2 (220-270 ms), P3 (300-400 ms), LPC1 (450-550 ms), and LPC2 (600-700). (B) Scalp topographies of the N2 and three late P3-like positivities depicted in (A) across task conditions (i.e., congruous, incongruous, and nogo).

### 3.3. RIDE analyses

Waveforms and scalp maps for the S, C, and R clusters are displayed in Figures 4 to 6, respectively. The S cluster appears to reflect mostly early sensory and attentional processes at stimulus onset. In the C cluster, the nogo condition elicits a large central P3-like positivity, whereas the incongruous condition elicits a flatter P3/LPC-like centro-parietal complex. In the R cluster, the positive parietal peaks are closely associated with response time, as expected, with an additional frontal peak elicited by the incongruous trials. A summary of the simple effects for BMC analyses at the Fz site is presented in Table 4.

**Figure 4.**
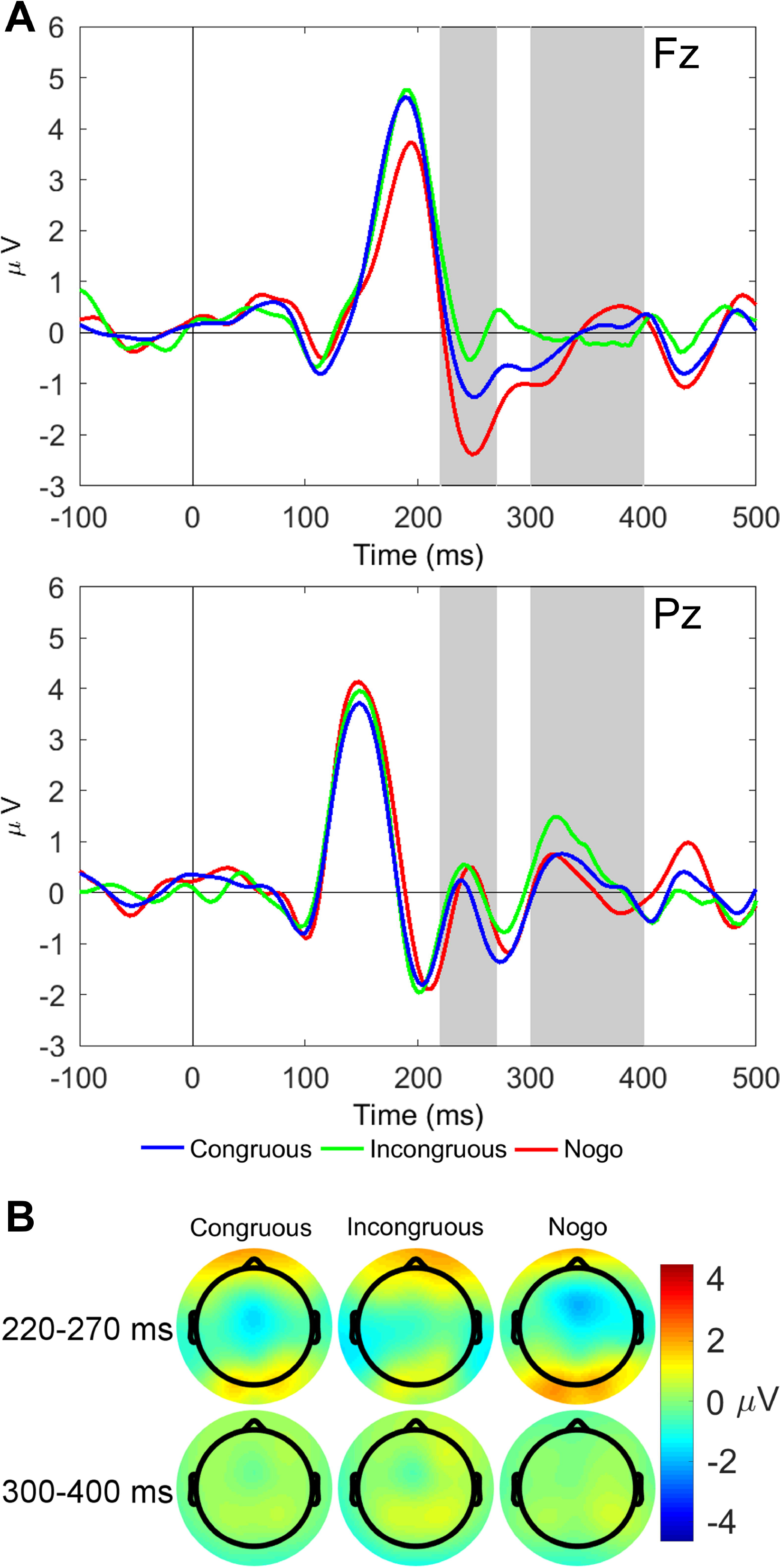
Stimulus-locked waveforms and scalp maps for the S cluster. (A) Waveforms depict grand-averages recorded from Fz (top) and Pz (bottom). The shaded area is the latency window used to measure P3-like activity in the S cluster: sN2 (220-270 ms), and sP3 (300-350 ms). (B) Scalp topographies for each task condition are mean amplitudes within the shaded time window in the waveforms.

**Figure 5.**
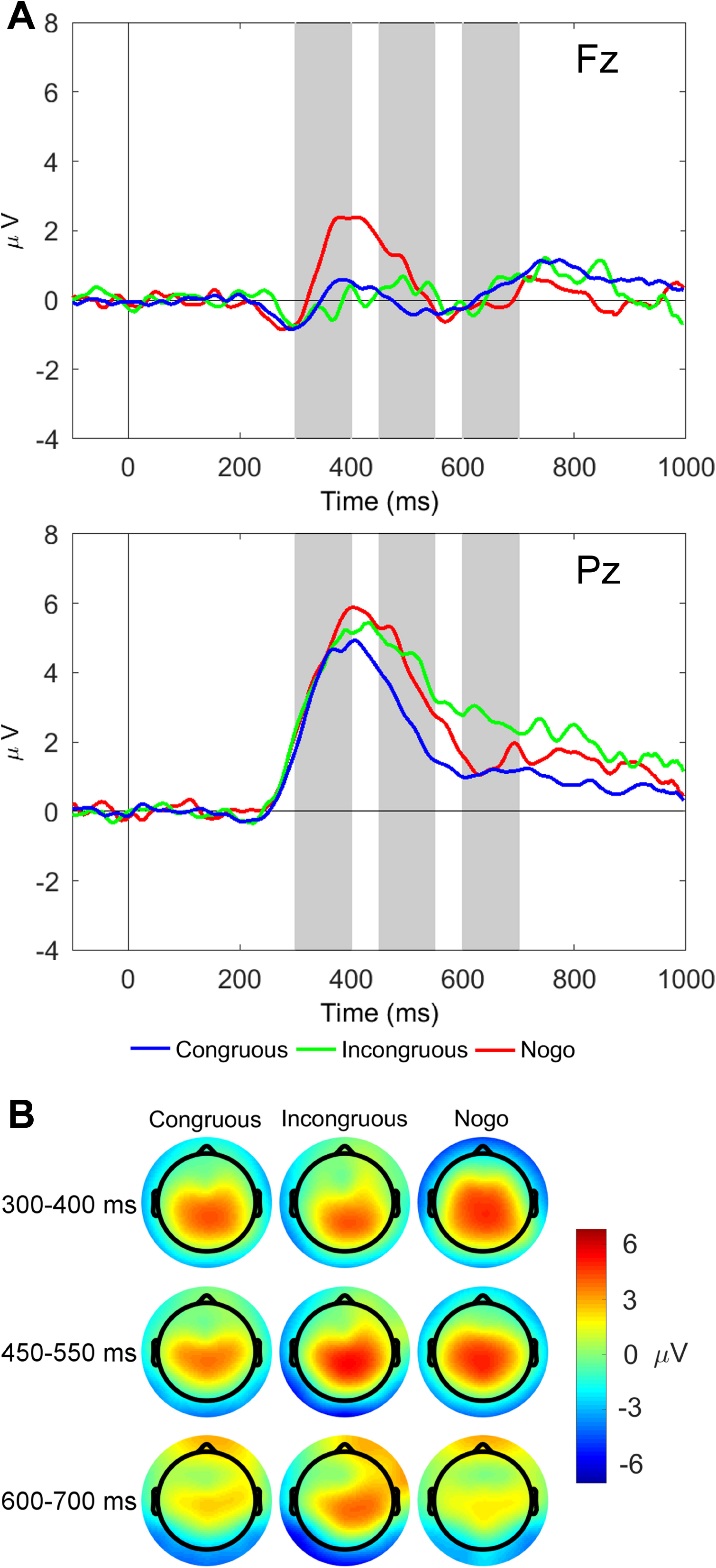
Latency-variable C cluster waveforms and scalp topography maps. (A) Waveforms depict mean voltages recorded from Fz (top) and Pz (bottom). Shaded areas indicate time windows used to measure mean amplitudes tracking the temporal dynamics of the C cluster P3-like complex: cP3 (300-400 ms), cLPC_1_ (450-550 ms), and cLPC_2_ (600-700 ms). (B) Scalp topographies of the three late cP3-like positivities depicted in (A) across task conditions (i.e., congruous, incongruous, and nogo).

**Figure 6.**
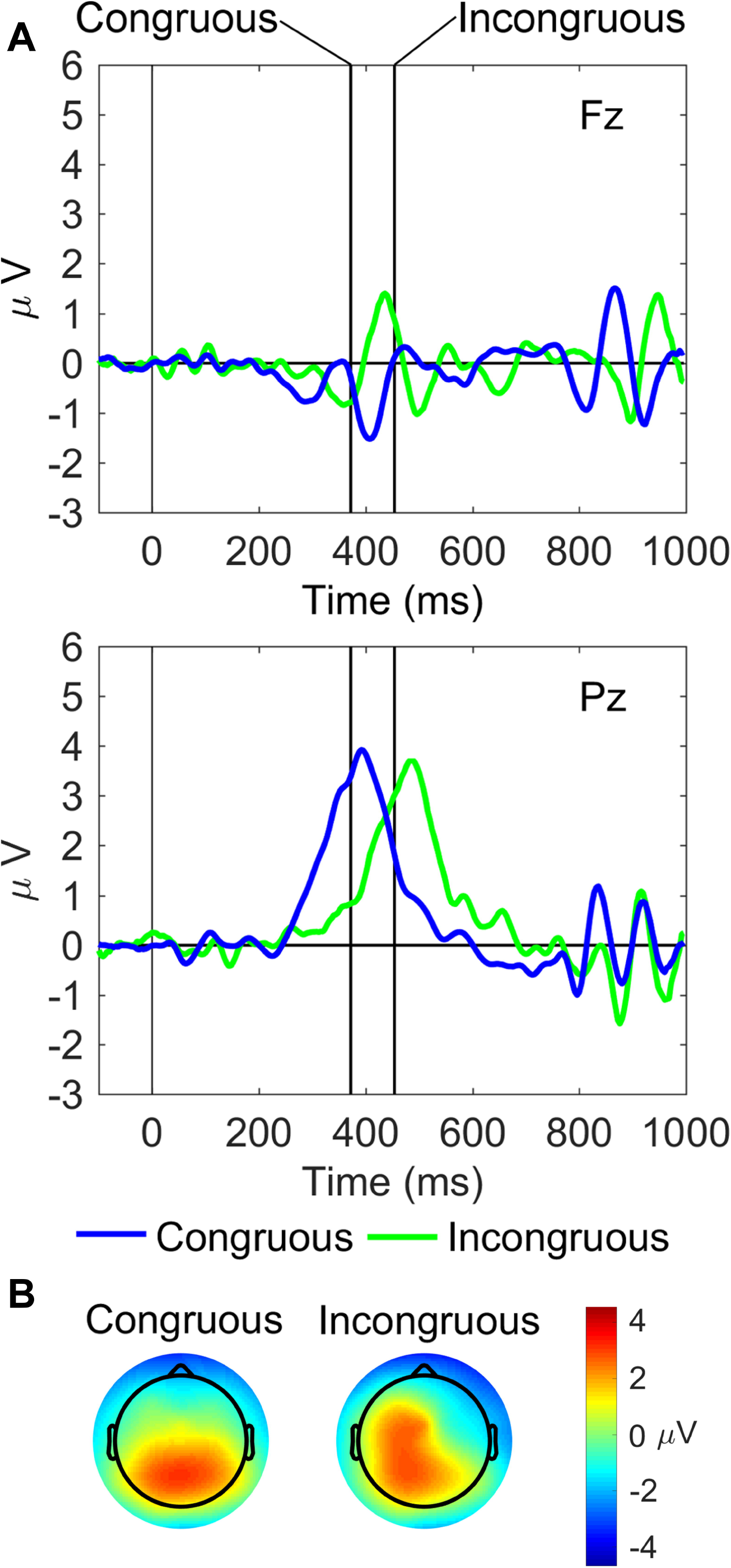
Response-locked waveforms and scalp maps for the R cluster. (A) Waveforms depict grand-averages recorded from Fz (top) and Pz (bottom). Vertical lines indicate the median response time for each task condition. (B) Scalp topographies for each task condition are the mean amplitudes measured in a 50 ms pre-response to 50 ms post response time window around the median response time for each condition.

#### 3.3.1. S cluster

For the sN2 component, the site × condition interaction and the main effect of condition were both significant (Table 3). Paired-samples t-tests found that mean sN2 amplitudes elicited by the nogo condition were larger (more negative) at Fz than at Pz (*t*(44) = 2.12, *p* = .040). BMC testing at Fz found evidence for differences between conditions (BF_10_ = 2987.04, *p*(H|D) > .99), revealing strong evidence for differences between all conditions (nogo > congruous > incongruous, where ‘>‘ indicates more negative; nogo-congruous BF_10_ = 6.42; nogo-incongruous BF_10_ = 295.30; congruous-incongruous BF_10_ = 10.54).

For the sP3 component, both main effects and the site × condition interaction were significant (Table 3). Paired-samples *t*-tests found that sP3 amplitude at Pz was larger in the incongruous condition than both in the congruous (*t*(44) = 2.49, *p* = .017) and the nogo conditions (*t*(44) = 3.37, *p* = .002). BMC testing at Fz found evidence against a difference between any conditions (BF_10_ = 0.07, *p*(H|D) = .07).

#### 3.3.2. C cluster

For the cP3 component, the site × condition interaction was not significant, whereas there were main effects of site and condition (see Table 3). Post-hoc tests showed that cP3 amplitude was maximal at Pz in comparison to Fz (*t*(44) = 6.42, *p* < .001), and the cP3 amplitude was larger for nogo than congruous and incongruous conditions (*t*(44) = 3.29, *p* = .002, and *t*(44) = 2.71, *p* = .010), respectively). BMC testing at Fz found evidence in favor of a difference between conditions (BF_10_ = 13.72, *p*(H|D) = .93), showing evidence for differences between nogo and the other two conditions (nogo-congruous BF_10_ = 11.53; nogo-incongruous BF_10_ = 4.65).

For the cLPC_1_ component, the site × condition interaction was not significant, but the main effects of site and condition both reached significance (Table 3). Maximal cLPC_1_ amplitude was found at Pz in comparison to Fz (*t*(44) = 6.02, *p* < .001), and the cLPC_1_ amplitude was larger for the incongruous and nogo stimuli than the congruous stimuli (*t*(44) = 3.76, *p* = .001, and *t*(44) = 3.05, *p* = .004), respectively). BMC testing at Fz found weak evidence against differences between conditions (BF_10_ = 0.70, *p*(H|D) = .41).

For the cLPC_2_ component, the site × condition interaction was not significant, with significant main effects for site and condition (Table 3). Paired-samples t-tests showed that the mean amplitude was larger at Pz than Fz (*t*(44) = 2.71, *p* = .010), and that the cLPC2 amplitude elicited by the incongruous stimuli was larger than the congruous stimuli (*t*(44) = 2.92, *p* = .006). BMC testing at Fz found evidence against differences between conditions (BF_10_ = 0.11, *p*(H|D) = .10).

#### 3.3.3. R cluster

The R cluster time window was 50 ms pre-response to 50 ms post-response for the congruous and incongruous conditions. The interaction and main effect of site reached significance (Table 3). Paired-samples t-tests found larger mean rP3 amplitudes in the incongruous than in the congruous condition at Fz (*t*(44) = 2.67, *p* = .011), while no such a difference was apparent at Pz (*t*(44) = 1.05, *p* = .30). BMC testing at Fz also found evidence in favor of a difference between congruous and incongruous conditions (BF_10_ = 4.26, *p*(H|D) = .81)

Finally, in light of the conspicuous differences in peak rP3 latency between conditions observed at Pz, a paired-samples t-test was conducted, which indicated that such differences were significant (*t*(44) = 8.93, *p* < .001, Cohen’s *d* = 1.33).

**Table 4.**
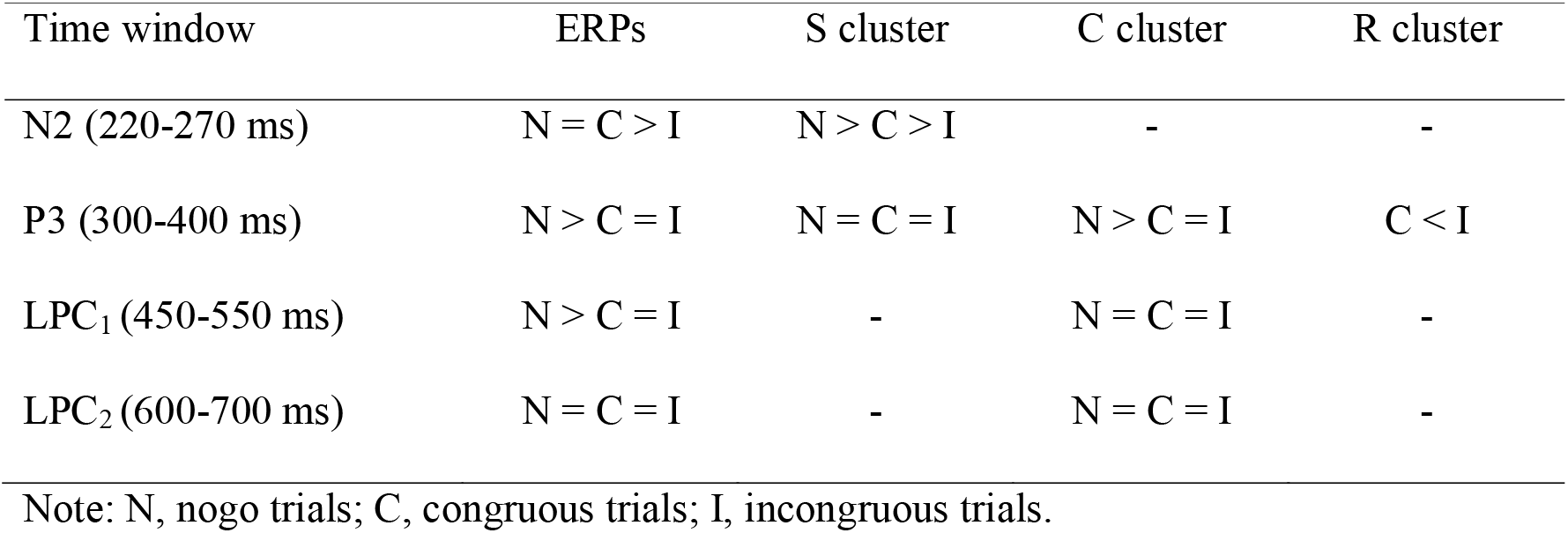
Summary of simple effects for Bayesian Model Comparisons at the Fz site.

| Time window | ERPs | S cluster | C cluster | R cluster |
| --- | --- | --- | --- | --- |
| N2 (220-270 ms) | $N = C > I$ | $N > C > I$ | - | - |
| P3 (300-400 ms) | $N > C = I$ | $N = C = I$ | $N > C = I$ | $C < I$ |
| LPC<sub>1</sub> (450-550 ms) | $N > C = I$ | - | $N = C = I$ | - |
| LPC<sub>2</sub> (600-700 ms) | $N = C = I$ | - | $N = C = I$ | - |

Note: N, nogo trials; C, congruous trials; I, incongruous trials.

### 3.4. Brain-behavior correlations

Lastly, Pearson product-moment linear correlations between RIDE decomposed amplitudes and behavioral measures (mean RTs, accuracy, and incongruency costs) were conducted. No correlations reached statistical significance (following Bonferroni correction). Additionally, in the light of the conspicuous differences in peak rP3 latency observed between congruous and incongruous trials, one final correlation was conducted between within-subjects differences in mean RT and the peak latency of the rP3 between those conditions. That correlation was statistically significant (*r* = .32, *p* = .033), with the scatterplot displaying a linear relationship (see Figure 7).

**Figure 7.**
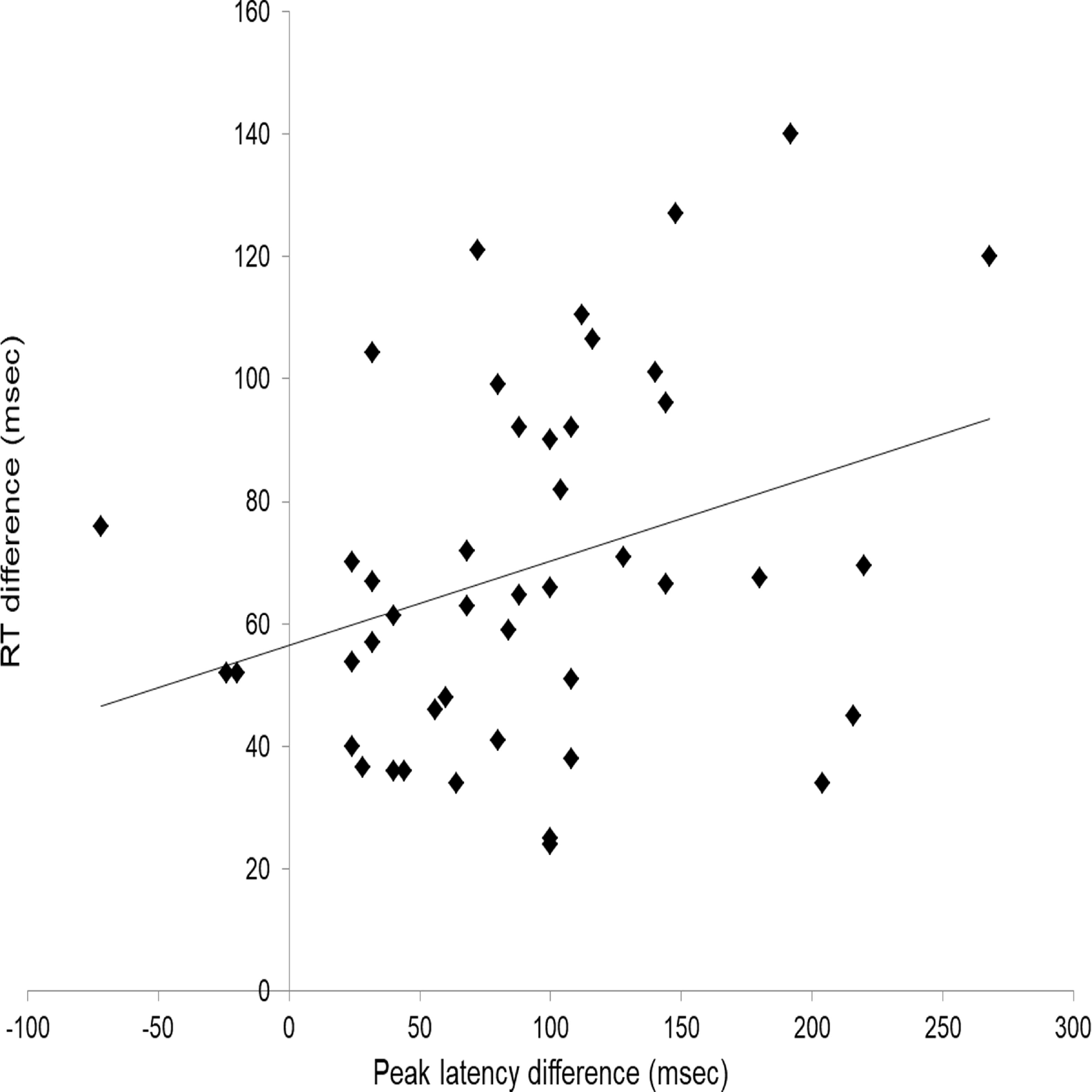
Scatterplot of within-subject difference in peak rP3 latency (msec; congruous subtracted from incongruous trials) and difference in mean reaction time (msec; congruous subtracted from incongruous trials).

## 4. Discussion

The current study aimed to expand upon Barceló and Cooper’s (2018) and Brydges and Barceló’s (2018) RIDE decomposition of target P3-like positivities during cognitive control of task switching, by examining these components in a flanker task. It was hypothesized that target (congruent and incongruent) and non-target (nogo) trials in the flanker task would elicit a N2-P3 complex indexing conflict detection and context updating operations that could be decomposed into stimulus-locked, intermediate, and response-locked sub-components. The mean amplitudes and scalp topographies of the N2-P3 complex were expected to be modulated by dynamic trial-by-trial adjustments in cognitive demands, with frontal effects predicted for those task conditions that overloaded working memory capacity (Fig. 1B; cf. Miller, 1956). More specifically, we examined whether one, two or three high-order task-set units (or latent variables) regulated performance in our flanker task in line with a well-known model of prefrontal executive control (Fig. 1A; Miller & Cohen, 2001). The results showed a highly dynamic picture of effects, with a stimulus-locked frontal N2 revealing early conflict detection and fast categorization of all three trial types, resulting in subsequent frontal P3-like activity (high-order context updating) elicited by infrequent nogo trials (C cluster) and by the perceptually difficult incongruous trials (R cluster). In turn, and as predicted by model 3 in Fig. 1B, the perceptually easier and more frequent congruous trials did not elicit frontal P3-like activity, although all three trial types did elicit parietally distributed P3-like activity (low-order context updating), mostly within the C cluster. These novel findings support the presence of up to three distinct high-order task-set units regulating performance in our flanker task (Fig. 1), as indexed by split-second, dynamic information transmission across frontal and parietal scalp regions, as is further discussed in more detail below.

Based on predictions from model 3 in Fig. 1, it was hypothesized that the nogo and incongruous trials of our flanker task would require larger degree of cognitive control and engage more prefrontal resources than the congruous condition, in line with the differential amounts of contextual information being transmitted by each of those trial conditions (Fig. 1B; Table 1). Under the assumption that dynamic trial-by-trial changes in cognitive demands associated with the processing of different flanker stimuli can be modelled as information transmission through hypothetical (hidden) high-order task-set units (Friston et al., 2017), we predicted differences on the electrophysiological indexes of conflict detection (frontal N2) and context updating (fronto-parietal P3) depending on the putative number of high-order task-set units regulating performance of this hybrid go/nogo flanker task (Fig. 1A,B; Table 1; see Supplementary materials). As such, it was expected that larger amplitudes of the frontal N2 and fronto-parietal P3 components will be associated with a larger number of high-order task-set units involved. Our conventional ERP results were generally consistent with previous studies, although these necessarily represent a compound mixture of different component processes. Thus, we found that the classic P3 was enhanced at Fz for nogo trials compared to congruous and incongruous trials, and the nogo-P3 was smaller over parietal regions (Fig. 3), in line with previous research (e.g., Gajewski & Falkenstein, 2013). Both nogo and incongruous trials elicited larger frontal LPC1 amplitudes than congruous trials, with incongruous trials eliciting significantly larger amplitude at Pz as well, consistent with previous research examining the LPC and task difficulty (Brydges, Fox, Reid, & Anderson, 2014). Now, in the light of our novel RIDE findings, these conventional target-P3 and nogo-P3 effects can be reinterpreted as a combination of stimulus-locked, response-locked, and latency-variable P3-like component processes.

Importantly, our novel RIDE results help clarify the functional significance of distinct component operations tracked by the frontal N2 and fronto-parietal P3-like modulations observed in all three clusters. In line with Brydges and Barceló’s (2018) findings, the largest P3-like modulations were captured by the C cluster, which is likely to be indicative of higher-order cognitive control operations, such as updating of high-order response policies given the sensory information and relative frequency of the incoming stimuli (i.e., frequent green fish vs. infrequent red fish). Interestingly, the S cluster captured not only early stimulus-locked peaks (i.e., a P2-like peak at both frontal and parietal sites between 140-200 ms), but also revealed later frontal N2 and parietal P3-like activity putatively indexing both high- and low-order transfer of information respectively from early stimulus-locked conflict detection to context updating triggered by changes in the sensory features of visual stimulation, since task condition also modulated the parietal sP3 (i.e., low-order context updating; see Fig. 4). From the context updating theory of the P3 (Donchin & Coles, 1998), this implies that changes in the perceptual aspects of the environment may trigger context updating at both high-order (Fz) and low-order (Pz) levels in the hierarchy of cognitive control, which was the case for our incongruous trials, as predicted by model 3 in Fig. 1B. There was also a visible response-locked P3-like component (rP3) with maximal parietal scalp distribution, likely reflecting “context updating” operations elicited by changes in motor or premotor control units associated with trial-by-trial variability in response selection. Importantly, and consistent with predictions from model 3, the more cognitively demanding incongruous condition elicited a significantly larger frontal rP3 and this peaked later over parietal regions than the easier congruous condition (cf. Brydges & Barceló, 2018). This suggests that the low-level SR mapping for congruous trials was easier to implement because it was more frequent and thus might have been naturally adopted as the default stimulus-response mode. Besides, all sensory information presented on the screen helped participants select the appropriate motor response, whereas in order to implement the correct S-R mapping for incongruous trials, participants must effortfully select just the contextually isolated central target and ignore all surrounding flankers, which presumably delayed the peak latency of parietal rP3 component downstream. Overall, these results support the findings of Brydges and Barceló (2018), in that changes in sensory, motor and sensorimotor levels of representation in the hierarchy of cognitive control can all trigger context updating mechanisms that differentially engage fronto-parietal regions from 220 to 700 ms post-stimulus onset and beyond, possibly reflecting either individual differences in the activation of various nodes across the fronto parietal cortical network for cognitive control (Bledowski et al., 2004; Duncan, 2010), or else temporarily recurrent fronto-parietal activation in those task conditions engaging larger cognitive demands (i.e., incongruous trials).

In the S cluster, frontal sN2 amplitude was modulated by task condition (nogo > congruous > incongruous; with nogo being most negative). The BMC analyses provided evidence of differences in frontal sN2 amplitude between all three conditions, implying that conflict detection is a stimulus-locked process that momentarily involved three distinct high-order latent task set units being held at prefrontal cortices (cf. model 3 in Fig. 1B), and has been putatively associated to the anterior cingulate cortex (Botvinick et al., 2004; Carter & van Veen, 2007; van Veen & Carter, 2002). This could be because it takes more perceptual resources (and time) to identify a perceptually difficult incongruous stimulus in comparison to a perceptually easy congruous stimulus (i.e., to identify one odd fish among the five exemplars in the display). Additionally, this is more time consuming and error prone because incongruous trials were less frequent (and hence more surprising) than congruous trials. Conversely, the absence of a frontal sP3 for any task condition provided strong evidence about only one high-order task set unit remaining active at this later latency window (cf. model 1; Fig. 1B). In spite of an absent frontal sP3 for any task condition, larger parietal sP3 amplitudes were elicited by incongruous trials than congruous and nogo trials. This finding suggests that low-order context updating may be triggered by a relatively infrequent and sensory complex visual target that differentially engages temporo-parietal regions of the fronto-parietal network under conditions of increased cognitive demands. Thus, low-order context updating triggered by relatively surprising perceptual changes in the environment enhanced parietal sP3 amplitudes on incongruous trials. This evidence suggests that this specific type of perceptually triggered context updating recruited more resources over temporo-parietal cortical regions in response to incongruous trials than to congruous and nogo trials. This finding suggests a highly dynamic and context-sensitive functioning of fronto-parietal networks, with fast split-second fluctuations in the amount of frontal and parietal resources that need to be engaged for processing each trial type (cf. Barceló & Cooper, 2018; Kieffaber & Hetrick, 2005). Additionally, the fact that the frontal N2 component elicited by the nogo condition was best captured by the S cluster rather than the C cluster suggests that what is traditionally considered to be the frontal nogo-N2 may largely reflect conflict detection triggered by stimulus-locked information transmission across prefrontal cortices (Carter & Van Veen, 2007; Nieuwenhuis et al., 2004; Van Veen & Carter, 2002). Specifically, the neural output of the frontal N2 may feed forward onto further high-order context updating operations (frontal P3), and even further down in the hierarchy of control into response inhibition and response selection operations at posterior association cortices (i.e., low-order context-updating; Fig. 1A), which could thus explain the modulations of subsequent processing stages captured by the latency variable and response-locked P3-like positivities (Botvinick et al., 2004; Gajewski, Stoerig, & Falkenstein, 2008).

In the central cluster, the cP3 peak (300-400 ms; the typical latency of classic P3 potentials) was modulated by high-order context updating, implying that the latency-variable cP3 was associated with higher-order updating of sensorimotor rules. Specifically, the BMC analyses on the cP3 component revealed the transient engagement of two high-order task set units, one for updating to infrequent nogo S-R links and one for updating to frequent target SR links (i.e., thus supporting model 2 in Fig. 1B; Table 5). This was consistent with the frontal nogo cP3 to relatively infrequent nogo trials requiring upholding a response, together with the absence of any frontal cP3 activity to the relatively frequent target trials requiring a motor response (congruous and incongruous trials). In contrast, the cLPC_1_ and cLPC_2_ were not modulated by high-order context updating (i.e., there was moderate BMC evidence in favor of only one high-order task set unit from Fz data, therefore supporting model 1 in Fig. 1B). In contrast, both cLPC_1_ components reached maximal amplitudes over parietal regions, where nogo and incongruous trials elicited greater amplitudes than congruous trials, whereas incongruous trials continued eliciting greater amplitudes than congruous trials in the cLPC2 time window. This pattern of results suggests that cognitive control is a fast and highly dynamic process, whereby once conflict is detected at frontal cortical regions (i.e., indexed by the frontal sN2 component), then context updating proceeds, first at high-order frontal regions (i.e., the nogo-cP3), and then later at low-order temporo-parietal association cortices (cLPC_1_, cLPC_2_) to update and reconfigure the S-R mappings needed to adaptively deal with the different task conditions. These findings suggest that the latency-variable components elicited in the C cluster capture several functionally distinct time-varying cognitive control operations resulting in subtly different P3-like scalp topographies as required by dynamical trial-by-trial changes in cognitive demands (cf. Barceló & Cooper, 2018; Brydges & Barceló, 2018).

In the response-locked cluster, a parietal P3-like positivity (rP3) showed a similar mean amplitude for congruous and incongruous trials, attesting for similar low-level context updating operations in both trial conditions. Most interestingly, though, there was an 80 ms difference in peak rP3 amplitudes between conditions, suggesting that low-level context updating for incongruous trials was delayed regarding the comparatively easier congruous trials. This difference in parietal peak rP3 amplitudes significantly correlated with behavioral efficiency, and is also consistent with predictions from model 3 in Fig. 1B, whereby only the highly informative and perceptually difficult incongruous trials elicited a frontal rP3 (high-order context updating) that influenced subsequent processes downstream by delaying the parietal rP3, an index of low-order S-R remapping (low-order context updating). Note that these more nuanced context updating operations are all response-locked and would thus remain hidden in traditional ERP waveforms (Fig. 3). This finding is similar to that reported by Brydges and Barceló (2018), whereby only the most cognitively demanding target trials immediately following a switch cue elicited additional frontal rP3 positivities, in spite of using identical visual displays as in target trials following a repeat cue. Brydges and Barceló (2018) argued that the extra cognitive demands required to categorize first target trials after a switch cue were contingent to response demands, and thus linked to the updating of low-order S-R mappings in a particular target trial. The present rP3 results are consistent with the increased response conflict in incongruous flanker trials, whereby the updating of low-level S-R mappings arguably involves concurrent selection and suppression of appropriate and inappropriate S-R links, respectively (i.e., stopping a default congruous response policy and adopt the reverse incongruous response policy; cf. Friston, et al., 2017). Thus, participants may adopt a default (or ‘prior’) congruous green-go response policy to maximize their behavioral efficiency in the most frequent congruous trials. However, when confronted with a relatively surprising incongruous green flanker stimulus, participants had to remap their low-level S-R links as part of the ongoing high-order task set unit, which results in a frontal rP3 followed by a delayed parietal rP3 in incongruous trials indexing the S-R remapping. Relatedly, Verleger, Grauhan, and Śmigasiewicz (2016) found that rP3 amplitude occurring at parietal sites approximately 40 ms pre-response was positively associated with task difficulty, and also that the most cognitively demanding trials (a rare response in a two-choice task) resulted in an additional fronto-central positivity occurring approximately 90 ms pre-response, thus generally consistent with the rP3 positivities observed in the current study.

The current study examined the phenomenon of the sustained fronto-parietal P3-like positivity elicited in cognitive control tasks such as task-switching and flanker tasks, and investigated the novel hypothesis from Brydges and Barceló (2018) that there are multiple stimulus-locked, latency variable, and response-locked peaks in the ERP that are modulated by task demands. Overall, the results from the RIDE analyses support this hypothesis. Supporting the results of Brydges and Barceló (2018), the present RIDE analyses show that the traditional dichotomy of the P300 complex (i.e., a frontal P3a and a parietal P3b subcomponent; Friedman et al., 2001; Polich, 2007) seems overly simplistic, given that the P3-like positivities elicited as a result of cognitive demands are far more nuanced than previously thought. The results also support Brydges and Barceló’s (2018) proposal that the target P300 complex comprises of a wide family of P3-like positivities overlying both frontal and parietal regions, possibly reflecting neural activation across a widely distributed fronto-parietal cortical network for cognitive control. These P3-like positivities are distinct from each other both functionally and anatomically, yet they appear to index a variety of context updating operations that require cognitive control, some of which are stimulus-locked, some response-locked, and some others that are latency-variable. These positivities overlay fronto-parietal scalp regions, with the amount of frontal recruitment depending on dynamic trial-by-trial changes in cognitive demands (i.e., working memory load; Miller, 1956), supporting Koechlin and Summerfield’s (2007) rostro-caudal axis of cognitive control and Miller and Cohen’s (2001) integrative theory of prefrontal function. In conclusion, the current study has shown that successful conflict detection and context updating are associated with a distinct combination of stimulus-locked, response-locked and latency variably electrophysiological processes putatively reflecting fast, split-second neural dynamics across a cingulo-fronto-parietal network for cognitive control (Botvinick et al., 2004; Duncan, 2010; Niendam et al., 2012).

## Acknowledgements

The research was funded by the School of Psychological Science at the University of Western Australia. Funding support was provided for by an Australian Postgraduate Award scholarship for Christopher Brydges. The funders had no role in study design, data collection and analysis, decision to publish, or preparation of the manuscript. The authors have declared that no competing interests exist.

